# Lack of mTORC2 signaling in CD11c^+^ myeloid cells inhibits their migration and ameliorates experimental colitis

**DOI:** 10.1101/2023.04.26.537895

**Authors:** Aline Ignacio, Marcella Cipelli, Tatiane Takiishi, Cristhiane Favero Aguiar, Fernanda Fernandes Terra, Bruno Ghirotto, Eloisa Martins Silva, Angela Castoldi, Yuli Thamires Magalhães, Tiago Antonio, Meire Ioshie Hiyane, Vinicius Andrade-Oliveira, Fabio Luis Forti, Niels Olsen Saraiva Camara

## Abstract

Mammalian target of rapamycin (mTOR) pathway plays a key role in determining immune cells function through modulation of their metabolic status. By specific deletion of Rictor in tissue-resident CD11c^+^ myeloid cells (CD11cRic^Δ/Δ^), this study investigated the role of mTOR complex 2 (mTORC2) signaling in dendritic cells (DCs) function in mice. We showed that upon DSS-induced colitis, lack of mTORC2 signaling CD11c^+^ cells diminish colonic inflammation, abrogates dendritic cell (DC) migration to the mesenteric lymph nodes (MLN), thereby diminishing the infiltration of T helper (Th) 17 cells in the lamina propria (LP). These findings corroborate with abrogation of cytoskeleton organization and decreased activation of Rac1 and Cdc42 GTPases observed in CD11c^+^-mTORC2-deficient cells. Meta-analysis on colonic samples from ulcerative colitis (UC) patients revealed increased gene expression of pro-inflammatory cytokines which coincided with augmented expression of mTOR pathway, positive correlation between the DC marker ITGAX, and IL-6, the expression of RICTOR, and CDC42. Together, this work proposes that targeting mTORC2 on DCs offers a key to hamper inflammatory responses and this way, ameliorates the progression and severity of intestinal inflammatory diseases.

## Introduction

Ulcerative colitis (UC) is the most prevalent inflammatory bowel disease (IBD) (1) which can lead to the development of colorectal cancer (CRC) (2). Along with genetic predisposition, microbial dysbiosis and lifestyle, the dysregulated production of pro-inflammatory cytokines drives the colonic inflammation and thus constitutes a promising target for therapeutic strategies. Indeed, the use of monoclonal anti-TNFα antibodies is an established treatment for IBD patients. However, the emerging number of patients that become unresponsive to the anti-tumor necrosis factor-α (TNFα) therapy (3, 4) highlights the importance of novel targets discovery for the improvement of current treatment approaches. Therefore, we sought to investigate other immune targets that could potentially be inhibited and served as an alternative to cytokines blockade.

In this context, dendritic cells (DCs) orchestrate the inflammatory immune response following UC onset by releasing pro-inflammatory cytokines and chemokines, which induce local innate immune cell activation and additional cell recruitment into the LP. The murine LP encompasses a heterogeneous group of conventional DC (cDC) identified as lineage (CD3, CD19, B220, NK1.1, CD64)-negative cells that express major histocompatibility complex (MHC) class II and the integrin CD11c. They can be divided in four subsets based on the expression of CD103 and CD11b expression: three major subsets (CD103^+^CD11b^-^, CD103^+^CD11b^+^ and CD103^-^CD11b^+^) and one minor (CD103^-^CD11b^-^) all able to migrate to MLN (5) and thus act as adaptive immune response initiators. In this environment, the cytokine milieu determines the functional characteristic of T cells where the prevalent differentiation of interleukin (IL)-17-producing T cells in the detrimental of regulatory T (Treg) cells is associated with UC pathogenesis (6, 7). Nevertheless, the mechanisms underlying the role of DCs in the initiation and development of IBDs is still not completely elucidated (8) where these cells can play a protective (9) and a detrimental (10) role on the disease.

Several studies point out to the importance of cell metabolism in determining the function and fate of immune cells. Mammalian target of rapamycin (mTOR) is a serine/threonine protein kinase that regulates cell proliferation, cell survival and protein synthesis in response to nutrients and mitogens. mTOR functions in two distinct complexes known as mTOR complex -1 (mTORC1) and -2 (mTORC2) that are distinguished by their accessory proteins, their sensitivity to the natural antibiotic rapamycin, as well as their effector functions. In contrast to mTORC1, prolonged rapamycin treatment is necessary for mTORC2 inhibition and its activators are still poorly defined. mTORC2 includes the proteins mTOR, mLST8, mSIN1 and the scaffolding protein Rictor, the latter essential for mTORC2 activity (11). Previous studies investigating the influence of mTORC2 on DCs responses showed that this complex regulates cytokine production and enhances their T cell stimulatory ability (12). However, little is known about its role in the function of tissue resident DCs that inhabit a complex environment, such as the colon. By using dextran sulfate sodium (DSS)-induced colitis model, and bone marrow-derived CD11c^+^ cells *in vitro* assays we conducted an exploratory assessment of the impact of mTORC2 absence in DC function upon acute inflammation.

## 1 Material and Methods

### 1.1 Mice

Male and female mice, 8-10 weeks old, were used for in vivo experiments. Female mice, 8-10 weeks old, were used for *in vitro* experiments. Rictor^flox/flox^ (Rictortm1.2Mgn) and CD11c-Cre (B6.Cg-Tg(Itgax-cre)1-1Reiz/J) mice were all on the C57BL/6 background and purchased from The Jackson Laboratory. We generated homozygous Rictor^flox/flox^ CD11c-Cre^+/+^ (CD11cRic^Δ/Δ^) or Rictor^flox/flox^ CD11c-Cre^-/-^ (Rictor^fl/fl^) by crossing Rictor^flox/flox^ with CD11c-Cre. Their genotypes were determined by PCR analysis of tail genomic DNA following The Jackson Laboratories protocols. Studies were performed on age and sex-matched littermates. All animal experiments were conducted in agreement with federal guidelines and approved by the institutional committee for animal care and use at the University of São Paulo (USP), Institute of Biomedical Sciences, São Paulo, Brazil. This study was registered under the protocol numbers 48/2014 and CEUA.048.2018.

### 1.2 Protein extraction and quantification

CD11c^+^ purified cells (1×10^6^) from bone marrow culture were lysed in Radioimmunoprecipitation assay (RIPA) buffer. Distal colon sections were homogenized on Tissue Lyser in 500 μL of RIPA followed by centrifugation at maximum speed at 4°C, for 10 minutes. Total protein quantification was determined using Pierce™ BCA Protein Assay Kit (Thermo Scientific Pierce™ BCA Protein Assay Kit) and samples were stored at -80°C until use.

### 1.3 Purification of CD11c^+^ cells

CD11c^+^ cells were isolated from spleen and bone marrow cultures using EasySep Mouse CD11c Positive Selection kit (Stem Cell Technologies) according to manufacturer’s instructions. Deletion of Rictor encoding gene was verified on cells isolated from the spleen by conventional PCR according to The Jackson Laboratories protocols for genotyping. mTOR pathway protein analysis was performed on cells purified from BM cultures.

### 1.4 Western blotting

Approximately 100 μg of total protein were prepared, separated by electrophoresis and transferred to nitrocellulose membrane accordingly.1 The molecular mass of protein was determined by comparison with the migration pattern of Precision Plus Protein™ Prestained Standards Dual Color (Biorad Laboratories, CA, USA). The primary antibodies used were anti-pAKT (S473), -AKT (9272), -pAKT (T308), -pS6 (T389) (1:1.000, Cell Signaling Technology, MA, USA), -Claudin 2 (1:500), -pJNK (1:500, Santa Cruz Biotechnology Inc), -β-actin (1: 10.000, Sigma-Aldrich, USA).

### 1.5 Colitis model

Colitis was induced by adding 3% DSS (36,000–50,000 MW, MP Biomedicals, Santa Ana, CA, USA) to the drinking water for 7 days ad libitum, changed on day 3. Control mice received water without DSS. After 7 days, animals were assessed by a clinical score as previously described (16) and euthanized for tissue harvest.

### 1.6 Histological analysis

Colon sections were formalin-fixed (10%) embedded in paraffin, deparaffinized and stained with hematoxylin-eosin (H&E) for histological analysis.

### 1.7 Enzyme-Linked Immunosorbent Assay

Enzyme-linked immunosorbent assays were used to quantify CRP, IL-6, IL-17, IL-23 and TNF-α from the intestinal tissue extract. Kits were purchased from R&D Systems (Minneapolis, MN, USA) and performed as described by the manufacturers.

### 1.8 Flow Cytometry

The colon was removed, opened longitudinally, washed with cold PBS (1x) to remove feces, and cut into 1 cm sections. After shaking into HBS solution (1x HBSS, 2% fetal calf serum inactivated (iFCS), 10 mM HEPES and 5 mM EDTA) at 37°C for 20 minutes, samples were washed 2-times with ice-cold 1x PBS. The tissue was minced prior to digestion in supplemented RPMI (Collagenase Type VIII (Sigma), DNAse I 10U/mL (Roche), 5% iFCS and 10 mM HEPES) at 37°C for 20 minutes. Digestion was deactivated using ice cold RPMI containing 10% iFCS and 10 mM HEPES at 1:1 ratio. The tissue was filtered through 100 um cell strainer, spun down and washed twice with FACS buffer (1x PBS, 2% iFCS, 2 mM EDTA). The MLN were minced and digested in supplemented RPMI (Collagenase Type 1A (Sigma), DNAse I 10U/mL (Roche), 5% iFCS and 10 mM HEPES) at 37°C for 20 minutes. The following steps were the same as for colon preparation. After counting in hemocytometer cells were labeled with mouse monoclonal antibodies using the markers live/dead Aqua (Life Technologies, Carlsbad, CA, USA), -CD45, -B220, -CD3, -siglecF, - CD64, -Nkp46, -CD11c, -CD11b, -I-A/I-E, -CD40, -CD80, -CD86, -CD4, CD8 (BD Biosciences). For cytokine staining cells were incubated in the presence of Golgi Plug (1x, BD Bioscience) for 4 hours, fixed and permeabilized using a Cytofix Cytoperm kit (BD Bioscience) and incubated with mouse monoclonal antibody anti-IFNγ, -IL17A, -IL-6, -IL12p40, -TNF-α (BD Bioscence). For transcription factor analysis, cells were fixed and permeabilized using Transcription Factor kit (eBioscience, San Diego, CA, USA) and then stained for Foxp3. Cells were acquired on a FACSCanto II (BD Biosciences) and data analysis was performed using FlowJo 9.5.3 software (T Tree Star, San Carlo, CA, USA).

### 1.9 Fluorescence in situ hybridization (FISH)

A segment of the colon containing feces was removed, fixed in methacarn (60% methanol, 30% chloroform, and 10% glacial acetic acid), embedded in paraffin, and cut into 5-μm sections. After deparaffinization, the sections were processed as recently published (43). The images were obtained using a confocal microscope, at a magnification of 40X.

### 1.10 Stool DNA extraction

Total DNA from fecal pellets was extracted using QIAmp DNA Stool Minikit (Qiage, Hilden, Germany) according to the manufacturer’s instructions. The DNA was evaluated for concentration and integrity and quantified by spectrophotometry (NanoDrop 2000, Thermo Scientific, Waltham, MA, USA).

### 1.11 16S Sequencing

The 16S rRNA gene was amplified using primer pairs selected from Klindworth23 targeting the V3 and V4 hypervariable region and 341F (CCTACGGGRSGCAGCAG) and 806R (GGACTACHV GGGTWTCTAAT). Amplicons were generated according to the Illumina MiSeq 16S Metagenomic Sequencing Library Preparation protocol and multiplexed using the Nextera XT Index kit. The quality of the libraries was assessed using Bioanalyzer (Agilent, Santa Clara, CA, USA) and quantified using Qubit (Life Technologies). Prepared libraries were pooled and then sequenced in a paired-end 2X 300-bp format on an Illumina MiSeq platform (San Diego, CA, USA). Using Dada2 2, phyloseq 3 and reshape2 packages 4 within R the forward and reverse reads were trimmed to 230 and 210 bp, respectively, and chimeras were removed using the remove Bimera Denovo function. Taxonomy was assigned using custom databases containing the 16S rRNA gene sequences of all the bacterial species used in the gnotobiotic models. Amplicon sequence variants (ASVs) that were present at <0.5% were excluded.

### 1.12 Bone marrow-derived CD11c^+^ cells generation

Bones were harvested and flushed with sterile 1x PBS. Cells were cultivated in IMDM (10% iFCS, 100 U ml–1 penicillin–streptomycin) with 20 ng/mL GM-CSF. On day 7, CD11c^+^ were purified and either plated with no further supplementation or stimulated with 100 ng/mL or 1 μg/mL of lipopolysaccharide (LPS, E. coli O111:B4, Sigma-Aldrich).

### 1.13 CD11c^+^-T cell co-culture

Purified CD11c^+^ myeloid cells from bone marrow culture were pulsed with 1 mg/mL of ovalbumin (Grade VI, Sigma-Aldrich) for 24 h. Splenic naïve OT II-CD4^+^ T cells were isolated using Mouse EasySep Naïve CD4^+^ T cell Isolation kit (Stem Cell Technologies) according to manufacturer’s instructions, and 1:10 CD11c^+^: T cell ration were cultured. For proliferation assay, CD4^+^ T cells were stained with Cell Trace (Life Technologies) and harvested on day 4. For the differentiation assay, cells were stimulated with stimulated with PMA (50 ng/mL) and ionomycin (750 ng/mL) in the presence of Stop Golgi 1x (Monensin, BD Bioscience), for 6 h. Cells were stained, acquired and analyzed as described previously.

### 1.14 Immunostaining of F-actin filaments

Purified CD11c^+^ myeloid cells (5 × 10^4^ cells) from bone marrow cultures were plated on an 8 well glass chamber slide (Nunc Lab-Tek-Thermo Fisher) in supplemented IMDM medium and incubated for 24 h, at 37°C. The medium was changed to contain any further additions or with 1 μg/mL of LPS and incubated for 1 or 2h, at 37°C. Cells were fixed in 4% PFA for 15 min following permeabilization cells 0.1% Triton-X (Sigma, Aldrich) for 15 min. Cells were washed and incubated overnight with Phalloidin (1:100) (Thermo Fisher, USA) and Hoechst (1:500) (Thermo Fisher, USA) in 1x PBS-2% BSA. Slides were mounted with VectaShield and analyzed in a Zeiss LSM 780-NLO confocal microscope (Carls Zeiss, Germany).

### 1.15 Chemotaxis assay

Purified CD11c^+^ myeloid cells (2 × 10^5^ cells) stained with mouse anti-CD11c antibody (BD Bioscience) were seeded on top of 8μM pore diameter polycarbonate inserts (Transwell system, Corning, USA) in supplemented IMDM medium (2% iFCS, 100 U ml–1 penicillin–streptomycin). Supplemented IMDM medium containing or not 200 ng/mL of murine CCL-19 (Cerdelane, Canada) were added in the bottom of the wells. The cells were stimulated with 1 μg/mL of LPS and incubated for 2 h, at 37°C. After incubation, the medium in the bottom of the wells were harvested and the migrated cells counted on flow cytometer FACSCanto II (BD Biosciences).

### 1.16 Rac1 and Cdc42 GTPases activity assay

The production of Pak-binding domain (PBD-GST) fusion protein was performed as previously described (59). To measure the endogenous Rac1 and Cdc42 activity, in vitro assays of pull-down were performed following a well-established protocol described (60). Purified CD11c^+^ myeloid cells (2×10^6^ cells/well) previously stimulated or not with 1 μg/mL of LPS for 2 h were disrupted by the RIPA lysis buffer. Protein quantification was performed using the Bradford (Bio-Rad, Hercules, CA) colorimetric method. For activity assay Rac1 and Cdc42, 250 μg of protein were incubated with 25 μg of PBD-GST beads at 4 °C for 90 min. The samples were centrifuge washed three times with buffer B (50 mmol/L Tris-HCl, pH 7.2, 1% Triton X-100, 150 mmol/L NaCl, 10 mmol/L MgCl2, 10 μg/mL leupeptin and aprotinin, and 0.1 mmol/L PMSF) and collected via centrifugation at 3,000 rpm for 3 min at 4 °C. Rac1- and Cdc42-GTP bound to PBD-GST-Sepharose beads were resolved on a 13% SDS-PAGE. The proteins were transferred to nitrocellulose membranes and blocked with 5% non-fat milk in TBS-T (20 mM Tris pH 7.6; 137 mM NaCl; 0.1% Tween) for 30 min at room temperature, following incubation for 4 h at room temperature either using a monoclonal anti-Rac1 antibody (1:500, Santa Cruz Biotechnology), or incubated overnight at 4 °C using a polyclonal anti-Cdc42 antibody (1:500, Santa Cruz Biotechnology). The membranes were incubated with the fluorescent secondary antibodies IRDye 680CW or IRDye 800CW for 1 h and visualized using the Odyssey Infrared Image System (CLx model, LI-COR).

### 1.17 Meta-analysis

The Gene Expression Omnibus (GEO) repository containing transcriptome datasets from UC patients was manually curated (https://www.ncbi.nlm.nih.gov/geo/). Two datasets (GSE75214 and GSE87466) containing representative expression data from UC patients (n = 74 and 87, respectively) and controls (n = 11 and 21, respectively) were selected for analysis. Only subjects with active disease were considered for analysis. The Differentially Expressed Genes (DEGs, Supplementary table 1) were filtered (log2-fold change (FC) ≥1, adjusted p value < 0.05) (61) and pseudogenes were excluded. KEGG Pathway enrichment analysis was carried using the Metascape software (Supplementary Table 3) (61, 62). The heatmap was generated using the Morpheus software (https://software.broadinstitute.org/morpheus/) based on the log2FC values for each gene throughout all studies analyzed. Correlation analyses between ITGAX and RICTOR, IL6 and CDC42 were performed.

### 1.18 Statistical analysis

Statistical analysis was performed using GraphPad Prism® software (San Diego, CA, USA). Differences among groups were compared using Student’s t-test, one-way or two-way ANOVA with Bonferroni post-test, as indicated. Based on the Kolmogorov-Smirnov test for normal distribution we selected Pearson or Spearman test for the correlation analysis of genes, as indicated in the respective figure. The observed differences were considered significant when p < 0.05 (5%).

## 2 Results

### 2.1 Rictor absence in CD11c^+^ myeloid cells attenuate DSS-induced colitis score

To investigate the role of mTORC2 signaling pathway in DC, we first confirmed deletion of Rictor in CD11c^+^ isolated from mice deficient in mTORC2 (named as CD11cRic^Δ/Δ^) under homeostatic condition (Supplementary Figure 1A). Next, we evaluated activation of mTORC1/2 pathways by immunoblotting in CD11c^+^ cells purified from bone marrow (BM) cultures from wild type mice (named as CD11cRic^fl/fl^) and CD11cRic^Δ/Δ^ mice. Although there was a slight decrease of Akt phosphorylation on threonine 308 in CD11c^+^ cells from CD11cRic^Δ/Δ^ mice, we observed similar levels of S6K phosphorylation on both CD11cRic^fl/fl^ and CD11cRic^Δ/Δ^ cells (Supplementary Figure 1B). These suggest that Rictor deletion does not impact on mTORC1 activity (13). In agreement with our results (Supplementary Figure 1A-B) and others (12, 13), CD11c^+^ cells isolated from CD11cRic^Δ/Δ^ mice displayed marked reduction in the phosphorylation of Akt on serine 473, an effect which was not observed on CD11cRic^fl/fl^ CD11c^+^ cells. This indicates a specific inhibition of mTORC2 in CD11c^+^ myeloid cells from CD11cRic^Δ/Δ^ mice allowing us to proceed on the investigation of the role of this signaling pathway on DC function.

DCs have a crucial role in intestinal homeostasis and development of acute and chronic inflammation (14, 15). In our CD11cRic^Δ/Δ^ mice, DSS-induced colitis resulted in attenuated weight loss, colon shortening and disease score compared to CD11cRic^fl/fl^ (Figure 1A-C). Moreover, histological injury of the colon with loss of crypt and goblet cells were attenuated. Analysis showed less colonocyte vanishing (Figure 2D) in CD11cRic^Δ/Δ^ mice. Although the C-reactive protein (CRP) was still increased after DSS administration, slighted lower levels were observed in Rictor-deficient mice compared to wild type CD11cRic^fl/fl^ mice (Figure 1E). These findings showed that lack of mTORC2 in CD11c^+^ cells ameliorates colitis score.

**Figure 1.**
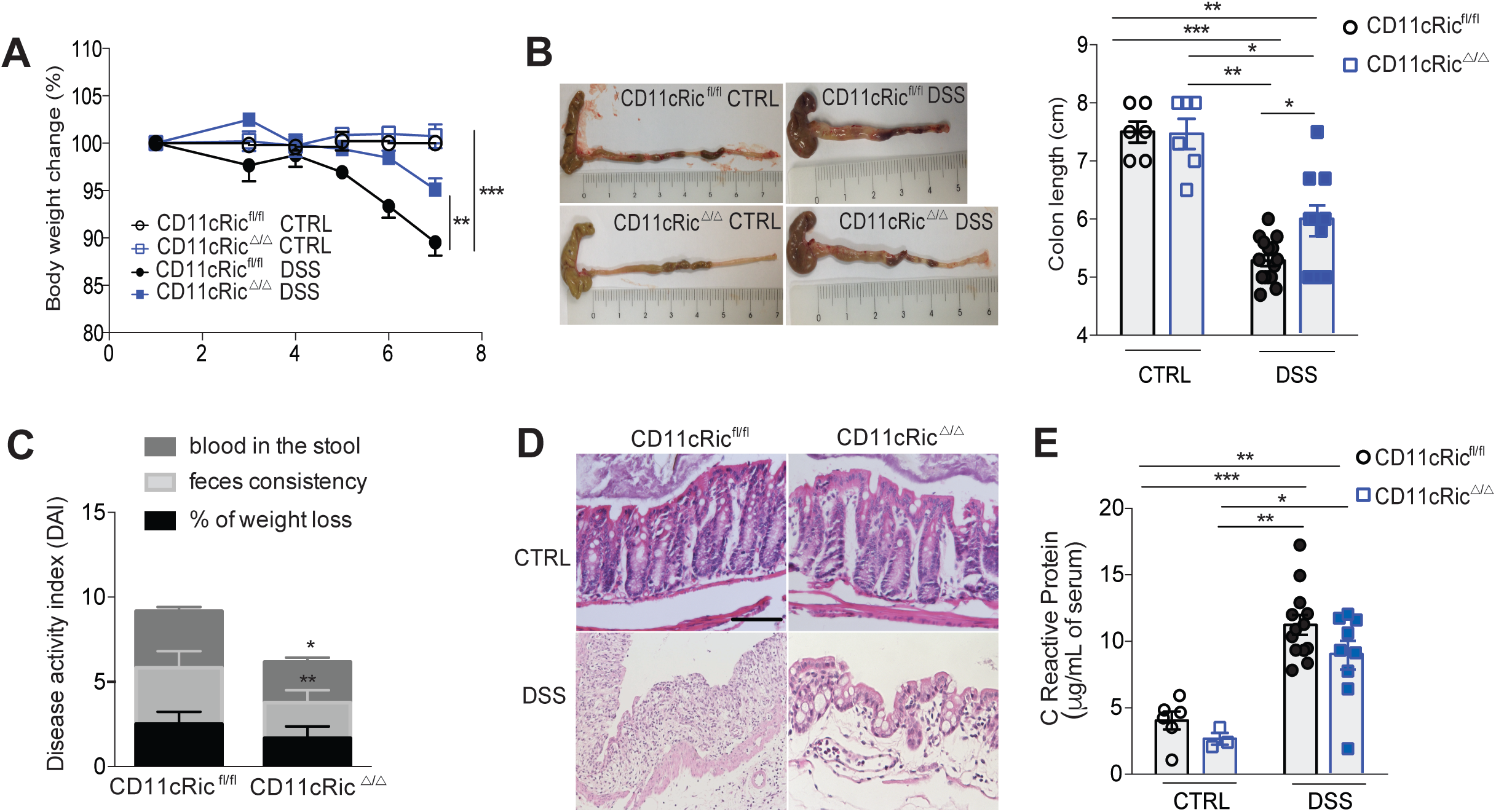
Lack of mTORC2 signaling in CD11c^+^ myeloid cells protects against colitis. Mice were treated with DSS 3% in the drinking water for 7 days. **A**, Body weight-change curves. **B**, Colon length. **C**, Colitis scores represented as Disease Activity Index (DAI). **D**, Representative histology of colon sections (scale bar 100 μm). **E**, C Reactive Protein in the serum. Data are from littermate controls CD11cRic^fl/fl^ (n = 5 – 13) and CD11cRic^Δ/Δ^ (n = 3 – 9) mice. Two-way ANOVA, Bonferroni post-test, *p < 0.05, **p < 0.01, ***p < 0.001; data are represented as mean ± SEM. Data are representative of three independent experiments.

**Figure 2.**
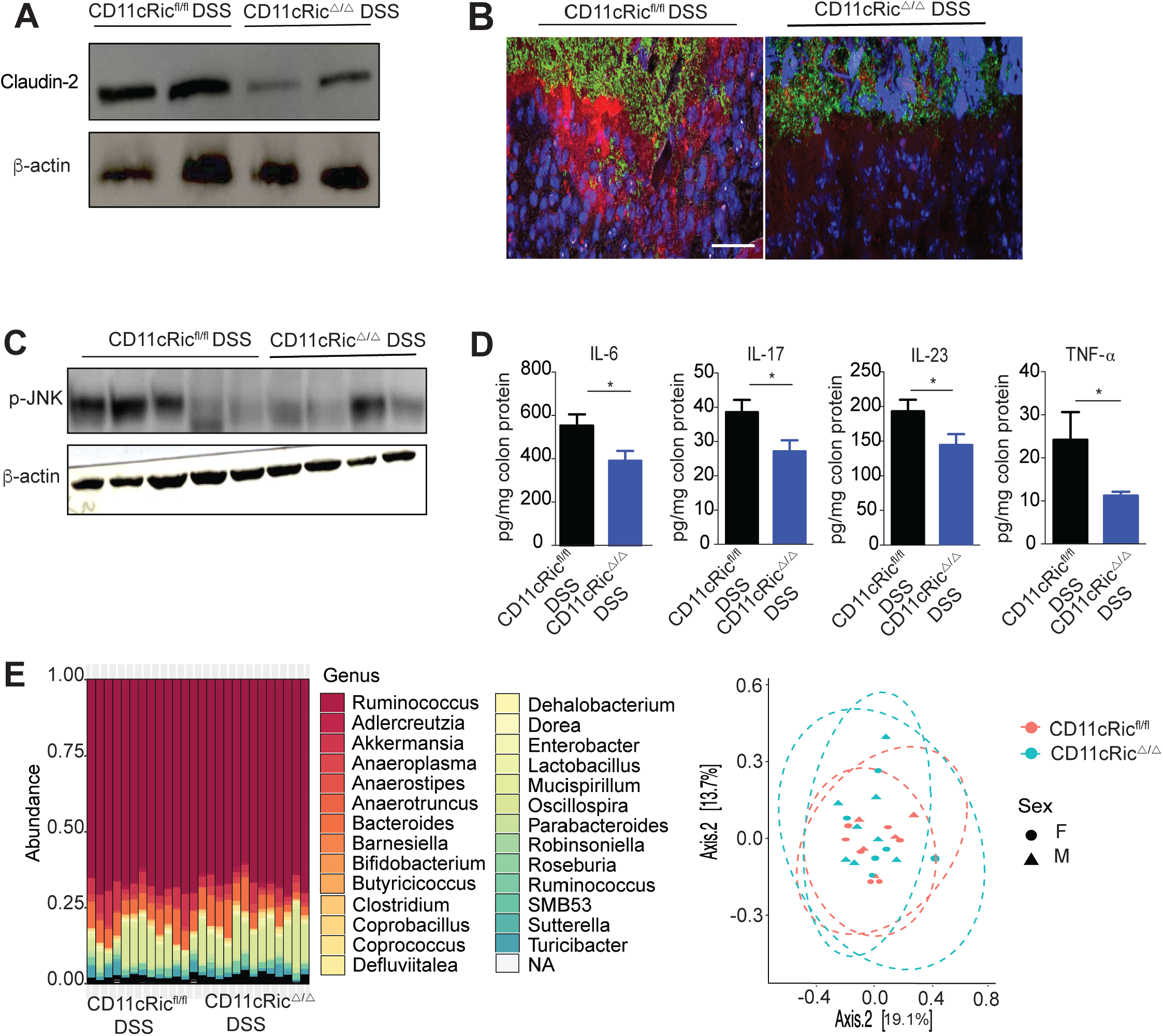
Epithelial barrier integrity and microbiota composition analysis from CD11cRic^fl/fl^ and CD11cRic^Δ/Δ^ mice. **A-D**, Mice were treated with DSS 3% in the drinking water for 7 days. **A**, Claudin-2 expression in the colon. **B**, Fluorescence in situ hybridization (FISH) analysis of total bacteria in the colon. Colon fragments containing feces were labeled with a probe for Eubacteria (green), epithelial cells nucleus were labeled with DAPI (blue) and mucus was labeled with Lectin-UEA1 (red) (scale bar 100 μm). **C**, Phosphorylated JNK in colon sections. **D**, Cytokine levels in colon segments. **e**, Microbiota composition analysis by 16S sequencing prior to DSS-induced colitis. Genera proportion (left) and Principal coordinate analysis (right). Data are representative of three independent experiments.

### 2.2 Decreased intestinal permeability and inflammation in CD11cRic^Δ/Δ^ is independent of gut microbiota composition

DSS model disrupts gut epithelial barrier with impairment of tight junction expression (16). In IBD, upregulation of Claudin-2 is associated with leaky gut and diarrhea (17, 18). We observed a reduction in the expression of Claudin-2 (Figure 2A) in CD11cRic^Δ/Δ^ mice. Those animals also showed less commensal bacteria proximity and invasion (Figure 2B) indicating preservation of intestinal barrier integrity.

In humans, activation of c-Jun N-terminal kinase (JNK) pathways in IBD patients has been shown to be higher in active sites of the inflammation (19, 20). In addition, it has been shown that the JNK pathway regulates T cell activation and the synthesis of pro-inflammatory cytokines (21, 22). CD11cRic^Δ/Δ^ mice showed less JNK activation (Figure 2C) along with reduced expression of the pro-inflammatory cytokines IL-6, IL-17, IL-23 and TNF-α in the colon (Figure 2D) upon colitis induction.

The colonic microbiota is the largest and most diverse bacterial community that inhabits the body surface (23). As changes in microbiota composition is an important factor that contributes to colitis development and severity in both humans (24) and mice (25), we wanted to evaluate whether lack of mTORC2 in CD11c^+^ cells in CD11cRic^Δ/Δ^ mice impact their intestinal microbiota composition. In order to avoid confounding factors such as maternal inheritance and long-term separating housing, (26) we harvested fecal pellets from CD11cRic^fl/fl^ and CD11cRic^Δ/Δ^ mice littermates-controlled mice prior to DSS treatment and performed 16S sequencing analysis. No differences were observed in the relative abundance (Figure 2E) and species diversity (Supplementary Figure 2B) between groups, indicating that microbiota composition does not play a role in attenuation of colitis score observed in CD11cRic^Δ/Δ^ mice.

### 2.3 CD11cRic^Δ/Δ^ shows DC migration impairment to MLN upon inflammation

Next, we evaluated the impact of mTORC2 deficiency on the abundance of DCs subsets within the lamina propria upon inflammation. The murine lamina propria encompasses four bona fide DC subsets (5). As shown in Figures 3A-B, we observed accumulation of all DCs subsets in the lamina propria of CD11cRic^Δ/Δ^ mice, mainly CD103^-^CD11b^-^ subset, with significant increase in numbers (Figure 3C). Two major DC populations could be identified in the MLN: the resident DCs (CD11c^+^MHCII^int^) and the migratory DCs (CD11c^+^MHCII^high^) (Figure 3D). All lamina propria DC subsets can migrate and can be found in the migratory compartment of MLNs and thus, have the potential to initiate adaptive immune responses (5). As we observed accumulation of DCs in the LP of CD11cRic^Δ/Δ^, we questioned whether this finding reflected a defect in DC migration to MLN. We observed a drastic reduction in the frequency and numbers of migratory DCs in CD11cRic^Δ/Δ^ (Figure 3E) with a significant decrease of all four DC subsets in the MLN (Figure 3F). Interestingly, CD11cRic^Δ/Δ^ also presented diminished DC frequency and numbers in the resident compartment (Supplementary Figure 3A), suggesting reduced input from BM following inflammation. It has been shown that expression of costimulatory molecules in bone marrow-derived DCs are altered by blocking mTORC2 signaling (12, 27). Given the importance of those molecules for DC function, we questioned whether drop in DC number in the MLN was also accompanied by impairment of costimulatory molecules expression. We observed no deficit in the expression of CD40, CD80 and CD86 for any DC subset in the MLN migratory compartment (Figure 3G), indicating that these cells can prime naïve T cells.

**Figure 3.**
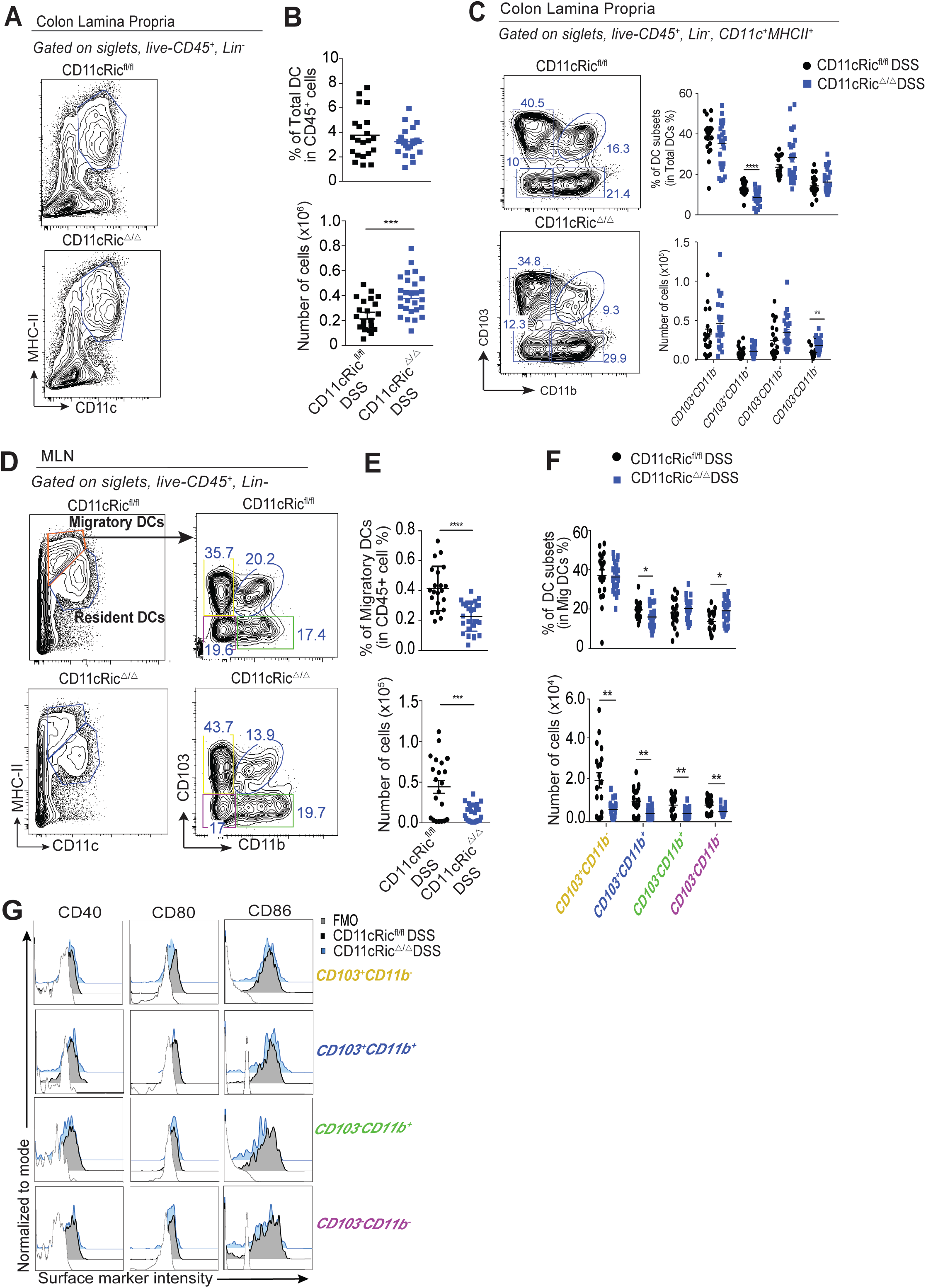
DCs in the colon Lamina Propria (cLP) and Mesenteric lymph nodes (MLN) from CD11cRic^fl/fl^ and CD11cRic^Δ/Δ^ mice. Mice were treated with DSS 3% in the drinking water for 7 days. **A**, Colonic DCs were identified as singlets, live, CD45^+^, Lin^-^ (CD3^-^, B220^-^, Ly6G^-^, SiglecF^-^, Nkp46^-^, CD64^-^) CD11c^+^MCHII^+^. **B**, Proportion and total cell counts of cLP-DCs. **C**, Proportion and total cell counts of DCs subsets in the cLP. **D**, Migratory DCs identified as singlets, live, CD45^+^, Lin^-^ (CD3^-^, B220^-^, Ly6G^-^, SiglecF^-^, Nkp46^-^, CD64^-^) CD11c^+^MCHII^high^, and respective subsets were evaluated in the MLN, **E. F**, Expression of costimulatory molecules CD40, CD80 and CD86 in migratory DCs subsets in the MLN. CD11cRic^fl/fl^ (n = 7 -8/experiment) and CD11cRic^Δ/Δ^ (n=8-13/experiment). Student T test*p < 0.05, **p < 0.01, ***p < 0.001. Data are expressed as mean ± SEM of three independent experiments pooled.

To understand whether the imbalance (accumulation in the LP and decrease in the MLN) on DC distribution in CD11cRic^Δ/Δ^ was a phenomenon linked to inflammation, we analyzed the lamina propria DC and migratory DC compartment in homeostasis condition. No changes between CD11cRic^fl/fl^ and CD11cRic^Δ/Δ^ were observed in the LP (Supplementary Figure 3B) and MLN (Supplementary Figure 3C-D) in a non-inflamed scenario, suggesting a possible delay in DC migration in CD11cRic^Δ/Δ^ after inflammatory trigger.

### 2.4 CD11cRic^Δ/Δ^ exhibited reduced Th17 and Treg in the lamina propria and MLN upon inflammation

It is known that IL-23-producing DCs promote the expansion of Th17 cells (28, 29). The excessive production of IL-17A by Th17 cells is a major player in colitis pathogenesis (30), where higher levels of mucosal IL-17A has been associated with higher rate of disease remission (4). As we observed reduced levels of IL-17 in CD11cRic^Δ/Δ^ lamina propria and lower numbers of DCs in the MLN, we hypothesized that Th17 differentiation was impaired when mTORC2 signaling is absent in CD11c^+^ cells. Although the total number of CD4^+^ T cells in the LP was not altered (Figure 4A), CD11cRic^Δ/Δ^ showed decreased numbers of IL-17^+^ CD4^+^ T cells (Figure 4B) and Tregs (Figure 4C) in the LP, with no impact in Th1 cells (Figure 4B). In agreement with our previous findings, reduced numbers of total CD4^+^ T cells were observed in the MLN of CD11cRic^Δ/Δ^ (Figure 4D), along with reduced IL-17-producing CD4^+^ T cells and Tregs (Figure 4E and F, respectively). We also observed a decrease in CD8^+^ T cell numbers in the LP of CD11cRic^Δ/Δ^ (Figure 4A), however with no changes in percentage of IL-17 and IFNγ production (Figure 4B). Reflecting the role of migratory DCs in the differentiation of naïve T cells, CD11cRic^Δ/Δ^ also showed reduced CD8^+^ T cells (Figure 4A & D) with impairment of CD8^+^ type 1 cytotoxic cells (Supplementary Figure 4B) differentiation in the MLN. These observations support our hypothesis that lack of mTORC2 signaling in lamina propria DCs delay their migration to MLN upon inflammation, reducing the number of all DC subsets capable of inducing adaptive immune responses in a secondary lymphoid organ, which in turn result in lower numbers of differentiated-effector T cells traveling to the lamina propria. Consequently, CD11cRic^Δ/Δ^ mice showed reduced inflammation following DSS treatment.

**Figure 4.**
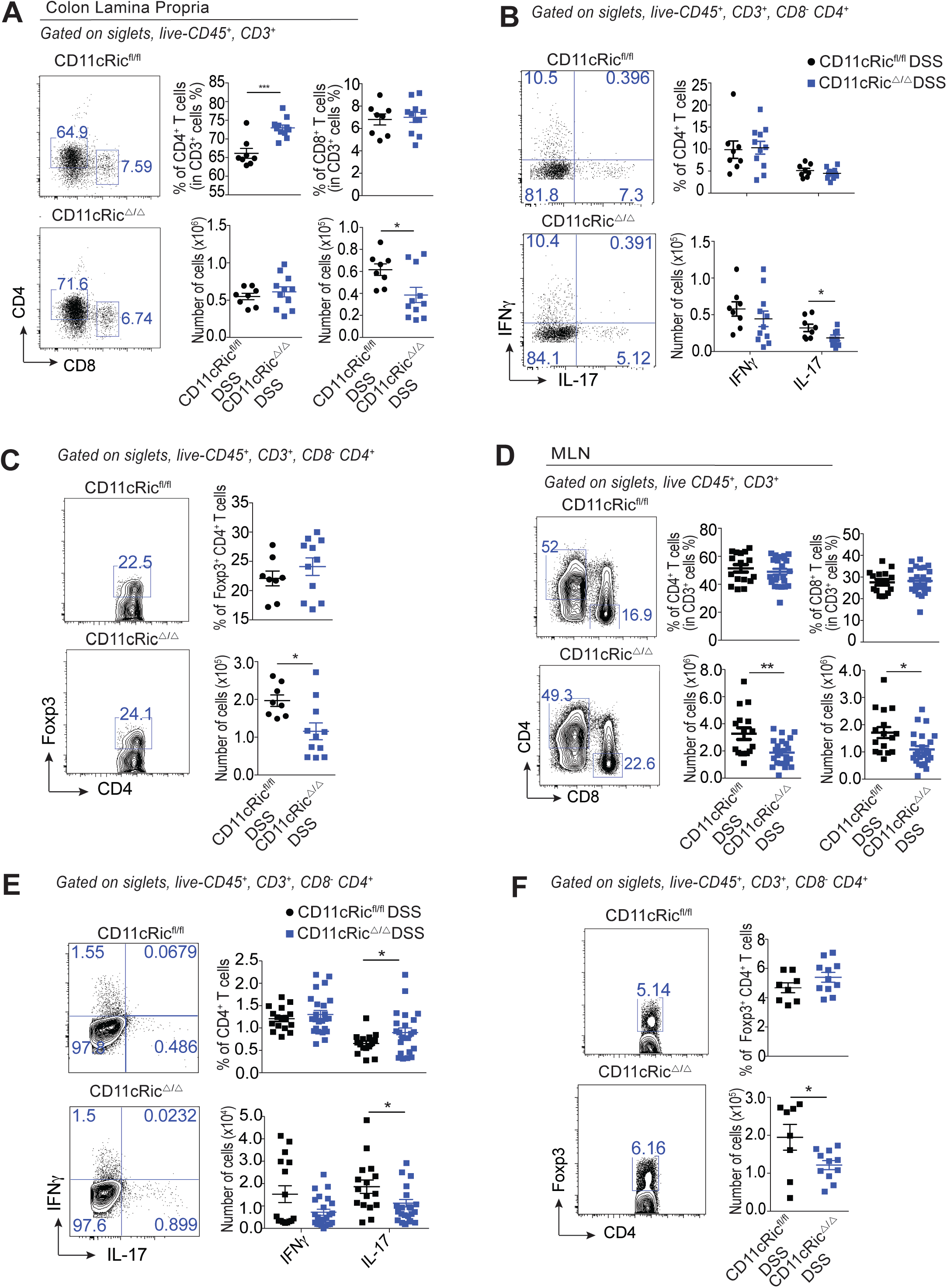
T cells in the colon Lamina Propria (cLP) and Mesenteric lymph nodes (MLN) from CD11cRic^fl/fl^ and CD11cRic^Δ/Δ^ mice. Mice were treated with DSS 3% in the drinking water for 7 days. **A**, CD4^+^ and CD8^+^ T cells in the cLP. **B**, IFN-*γ* and IL-17 production on CD4^+^ T cell from cLP. **C**, Tregs from cLP. **D**, CD4^+^ and CD8^+^ T cells in the MLN. **E**, IFN-*γ* and IL-17 production on CD4^+^ T cell from MLN. **F**, Tregs from MLN. CD11cRic^fl/fl^ (n = 7 -8 /experiment) and CD11cRic^Δ/Δ^ (n=5 - 6/experiment). Student T test *p < 0.05, **p < 0.01, ***p < 0.001. Data are expressed as mean ± SEM from two independent experiments pooled.

### 2.5 Rictor signaling regulates cytokine production and cytoskeleton organization in DCs

To further explore the mechanisms by which Rictor signaling regulates DCs migration, we performed *in vitro* assays using CD11c^+^ cells purified from GM-CSF bone marrow-derived myeloid cells culture (Supplementary Figure 5A). First, we evaluated the costimulatory molecules expression after stimulation with lipopolysaccharide (LPS) and similar to lamina propria DCs, CD11c^+^ cells from CD11cRic^Δ/Δ^ did not show impairment in the upregulation of CD40, CD80 and CD86 after inflammatory stimulus (Supplementary Figure 5B). It has been shown that lack of Rictor signaling in DCs alters pro-inflammatory cytokine production by those cells. (12, 27) Given that, we analyzed the cytokine production in LPS-stimulated CD11c^+^ cells and contrary to reported findings, lack of Rictor decreased the production of IL-6 and TNF-α, with no changes in IL-12/23p40 production (Figure 5A). As cytokine synthesis is determinant for the induction of T effector subsets, we co-cultured ovalbumin (OVA)-pulsed CD11c^+^ cells with naïve OT-II CD4+ T cells. CD11c^+^ myeloid cells from CD11cRic^Δ/Δ^ were able to induce proliferation of CD4^+^ T cells in higher proportions observed for CD11cRic^fl/fl^ (Supplementary Figure 5C). Next, we analyzed T cells subset differentiation in OVA-pulsed CD11c^+^ cell co-culture. As shown in Figure 5B, and consistent with our in vivo findings, OVA-pulsed CD11cRic^Δ/Δ^ decreased the frequency of IL-17-producing CD4+ T cells, without compromising interferon gamma (IFN*γ*)^+^CD4^+^ T cell differentiation. A rising trend in the incidence of Treg was noticed (Figure 9B). Together, these observations suggest that Rictor signaling partially modulates cytokine production in DCs which impacts differentiation of certain Th subsets.

**Figure 5.**
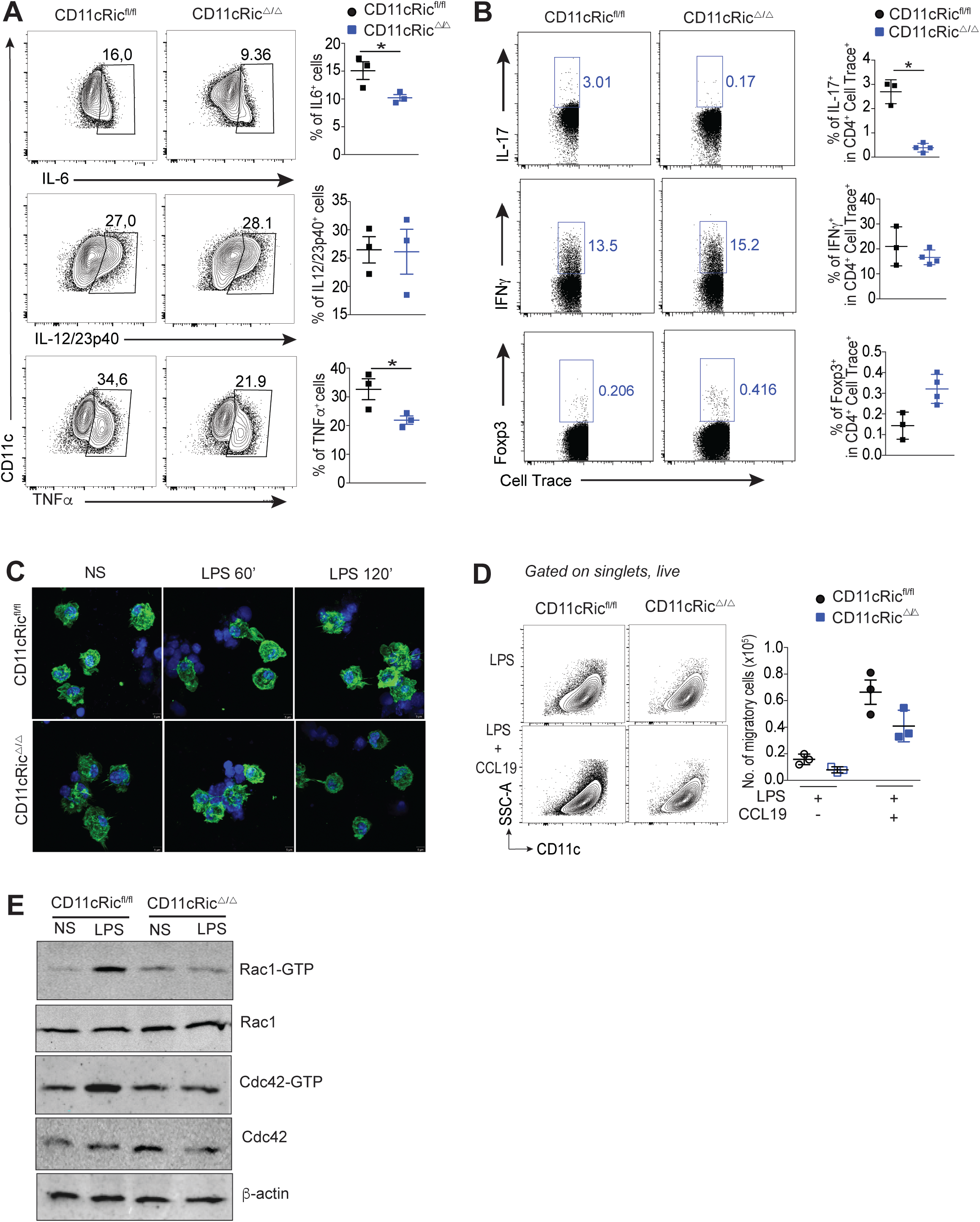
mTORC2 signaling modulates CD11c^+^ myeloid cells functions. **A**, GM-CSF differentiated bone marrow-derived CD11c^+^ from CD11cRic^fl/fl^ and CD11cRic^Δ/Δ^ mice were evaluated for the expression of pro-inflammatory cytokines following stimulation with LPS (100 ng/mL). **B**, T cell differentiation co-cultured with CD11c^+^ cells was evaluated on day 5. **B**, Polymerized F-actin was labeled with Phalloidin (green) and cell nucleus with Hoescht (blue) after stimulation with LPS (1μg/mL) for the indicated time points. **d**, Cell migration of LPS-stimulated (1μg/mL) CD11c^+^ cells under the presence or absence of the chemoattractant CCL19 (200 ng/mL). **e**, GTPases activity on unstimulated and LPS-stimulated (1μg/mL) CD11c^+^ cells. Student T test *p < 0.05. Data are expressed as mean ± SEM and representative from two or three independent experiments.

As activated DC increases cytoplasm membrane projection as result of cytoskeleton organization after activation, we stained polymerized F-actin filaments in unstimulated and LPS-stimulated cells. While similar shape was observed in unstimulated CD11c^+^ cells, shorter dendrites protrusion was noticed after 120 min in LPS-activated CD11cRic^Δ/Δ^ (Figure 5C). To assess their migratory capacity, chemotaxis of activated CD11c^+^ cells in the presence or absence of chemoattractant chemokine (C-C) motif) ligand 19 (CLL19) was analyzed. In CCL19 absence, few cells were able to migrate and no difference was observed between CD11cRic^fl/fl^ and CD11cRic^Δ/Δ^ However, in the presence of CCL19, migration of activated CD11c^+^ cells derived from CD11cRic^Δ/Δ^ mice was decreased (Figure 9D). Cytoskeleton organization and cell motility have been associated with the activity of Rho GTPase family, including RhoA, Rac1 and Cdc42 members (31, 32). Proper-cytoskeleton organization requires signaling through Rac1 and Cdc42, conferring to DCs their ability to promote efficient T cell activation (33). As mTORC2 function in cytoskeleton organization has been linked to cell migration in neutrophils (34) and cancer cells (35), we hypothesized that lack of Rictor abrogates GTPases activation and leads to impaired DC migration. LPS-activated CD11c^+^ cells derived from CD11cRic^Δ/Δ^ mice showed decreased activation of Rac1 and Cdc42 GTPases, with no changes in unstimulated cells (Figure 5E), indicating that mTORC2 plays a crucial role in DC migration machinery under an inflammatory condition.

To explore a possible translational impact of our findings, we analyzed the dataset of two cohorts of patients with UC (GSE75214 and GSE87466). The filtered DEGs (log2FC ≥ 1, adjusted p value < 0.05) in colonic biopsies from UC patients compared to healthy controls were selected (Supplementary Table 1). We observed that IL-17 and TNF-α signaling pathways are activated in UC biopsies (Figure 6A, Supplementary Table 2). Corroborating our results, among the genes found enriched, UC patients from both cohort studies showed increased expression of pro-inflammatory cytokines (IL-6, -12, -17A, -23A, TNFA and IFNG), the DC marker ITGAX, genes from mTOR pathway (RICTOR, AKT, FOXO1, S6K) and the Rho GTPases RAC1, RHOA and CDC42 (Figure 6B). We also observed a positive correlation between the expression of RICTOR and ITGAX, IL-6 and ITGAX and CDC42 and ITGAX in both datasets (Figure 6C) suggesting that in UC patients, mTORC2 signaling plays an active role in DC function, possibly by the regulation of cytokine production and DC migration.

**Figure 6.**
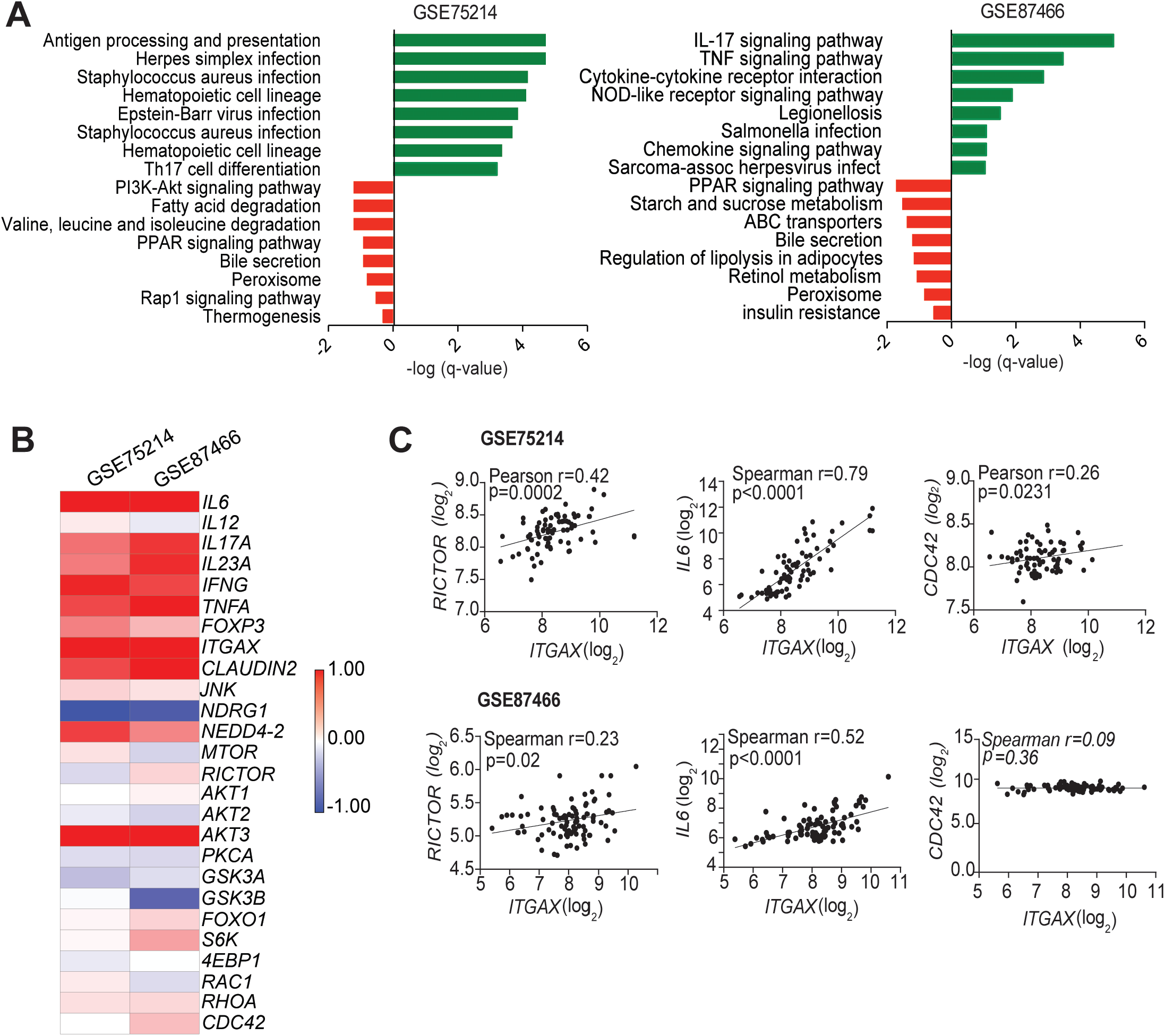
Meta-analysis in patients with UC. Expression of genes on colonic regions from UC and healthy control patients from two datasets (GSE75214 and GSE87466). **a**, Pathway enrichment analyses of down (logFC <1, p < 0.05) and upregulates (logFC > 1, p < 0.05) genes in UC patients from cohort GSE75214 (right) and GSE87466 (left). **b**, Immunological markers and mTOR molecules expressed in the colonic section from UC relative to healthy controls patients. **c**, Correlation analyses between the expression of ITGAX and -RICTOR, -IL6 and -CDC42 to evaluate the association between DCs and the upregulation of mTOR pathway, inflammatory immune response and cell migration, respectively, in UC patients.

## 3 Discussion

There is no doubt that mTOR is a central regulator of cellular metabolism, and therefore has a pivotal role on immunity and inflammation tone (36, 37, 38, 39). Nevertheless, little is known about its role in myeloid cells and it warrants further investigation on the biology of tissue resident cells. We demonstrated that lack of signaling through mTORC2 in CD11c^+^ cells diminishes the levels of mucosal pro-inflammatory cytokines, including IL-6, TNF-α, and IL-17, limits epithelial barrier damage, and reduces disease active index upon DSS-induced colitis. These findings can be explained by the decreased number of lamina propria migratory DCs observed in the mesenteric lymph nodes of mice mTORC2-deficient in CD11c^+^ cells, which resulted in little differentiation of effector T cells, particularly Th17. Treatment with the ATP-competitive mTOR inhibitor AZD8055 blocks the signaling pathway downstream to mTORC2 in several cell types, and it has been shown to attenuate intestinal inflammation by inhibiting Th17 infiltration into the colonic mucosa in the DSS-induced colitis model (40). Although this study supports our data, the use of AZD8055 by Hu et al. (40) may reflect an indirect effect in the decrease of Th17 cells, highlighting the importance of investigating how this pathway interferes in the function of individual cell types. Increased levels of IL-17 in the presence of augmented levels of IL-23 in IBD patients lead to activation of intestinal epithelial cells and fibroblasts that secrete IL-6, TNF-α and metalloproteinase, causing ulcerative lesions in the intestine wall (28, 41, 42). Similarly, we also showed attenuation of this scenario when mTORC2 signaling is impaired on LP-DCs cells. It is important to note that intestinal macrophages also express CD11c^+^ and therefore, CD11c-driven CRE (43). Although our study cannot rule out a possible influence of intestinal macrophages on the inflammatory response observed, a body of evidence suggests that mTORC2 activation in macrophages undergo a dramatic shift towards a proinflammatory phenotype. It has been reported that activated Rictor-deficient macrophages secrete increased levels of IL-6 and TNF-α (44). A worked performed in mice Rictor-deficient on macrophages showed that lack of mTORC2 signaling in these cells abrogates the differentiation of “wound-healing” M2 macrophages without affecting the differentiation of “inflammatory” M1 macrophages (45). Corroborating the opposite role of mTORC2 in macrophages, another study demonstrated that activation and not inhibition of mTORC2 in macrophages prevents aggravated disease of colitis-associated colorectal cancer in mice and humans (46). These studies support the evidence that amelioration on the disease activity observed in our study is most likely to be regulated by DCs mTORC2-deficient cells rather than macrophages.

In contrast to previous reports showing that lack of mTORC2 in stimulated-DCs induces overexpression of costimulatory molecules (16), here we showed that mTORC2-deficient LP-DCs did not overexpress MHC class II and the costimulatory molecules CD40, CD80 and CD86, after DSS treatment. Similar results were observed in our LPS-stimulated bone marrow-derived CD11c^+^ cells. A possible explanation for the divergence in the results is the influence of different microbiota composition on resident cells as mTOR is a sensor for environmental cues (36). It is important to note that we used the littermates not expressing cre recombinase rather than C57BL/6 wild type controls in all our experiments (30). In addition, the use of LPS derived from different bacteria in each study may be the cause of distinct cell responsiveness in the in vitro experiments. Although mTORC2 absence in CD11c^+^ cells did not impact their ability to induce CD4^+^ T cell proliferation in vitro, the polarization to Th17 was dampened in Rictor-deficient CD11c^+^ cells. Raiche-Regué et al. (16) have reported that Rictor deletion in DCs increases the production of pro-inflammatory cytokines and enhances their ability to expand Th1 and Th17 polarized cells. The discrepancy between findings may reflect different mouse lines used for knockdown mTORC2 activity. Whereas the latter authors used floxed Rictor mice crossed with C57BL/6 expressing Cre under the ROSA26 promoter, we used the constitutively Rictor knockout mice.

In the last two decades, 30% to 50% of IBD patients have shown loss of response to anti-TNFα monoclonal antibody therapy (47, 48, 49), prompting the scientific community to target an alternative mechanism in IBD. Controlling the early proinflammatory responses in IBD patients is crucial to limit tissue damage and the detrimental downstream cascade of dysregulated inflammatory events. The blockade of T-cell recruitment into lesions has been shown promising therapeutic results in humans (50, 51, 52) and thereby, place DCs in the center of this scenario as professional T-cell activators and mobilizers. We observed that lack of signaling through mTORC2 in CD11c+ cells inhibits actin cytoskeleton organization and migration. Supporting our findings, our group (53) and a recent study by Jangani (54) demonstrated that loss of mTORC2 activity impairs the migration of neutrophils and monocytes, respectively. Cytoskeleton organization and motility in humans and murine DCs have been classically associated with the Rho family GTPases (33, 40, 41). Defects in chemotaxis observed in mTORC2-deficient neutrophils has been also associated to Rho GTPases Rac and Cdc42 (34), indicating that Rac and Cdc42 act as downstream effector molecules of mTORC2 in actin cytoskeleton organization and migration of both in vivo and in vitro activated CD11c^+^ cells. To further investigate the relevance of our findings in the context of IBD in humans, we performed meta-analysis on cohorts of patients with UC compared with healthy controls. Our meta-analysis indicated that signaling pathways related to production of pro-inflammatory cytokines with concomitant activation of mTORC2 on CD11c^+^ cells were enriched in colonic biopsies from UC patients. These observations suggest that in humans, modulation of mTOR activity specifically in immune cells can also determine the progress of the disease. Finally, our findings indicate that specific mTOR inhibition could be a new target for UC treatment and shed a new light for mTOR in modulating DC function in humans and mice.

## Supporting information

Metanalysis

## 5 Conflict of Interest

*The authors declare that the research was conducted in the absence of any commercial or financial relationships that could be construed as a potential conflict of interest*.

## 6 Author Contributions

AI and NOSC conceived the project, interpreted the data and wrote the manuscript. AI designed and performed the experiments. TT participated in the experiments, data interpretation and reviewed the paper. MC and YTM performed and interpreted the GTPase assay. BG and EMS performed the meta-analysis. CFA, FFT, AC, VAO participated in the experiments. TA performed the 16S sequencing. MIY contributed to animal maintenance and genotyping. FLF supervised the GTPase assay and provided intellectual input on the paper. NOSC supervised the overall project.

## 7 Funding

This research was supported by Fundação de Amparo à Pesquisa do Estado de Paulo (FAPESP, Grant No 2014/05496-0 and 2017/05264-7). is study was financed in part by the Coordenação de Aperfeiçoamento de Pessoal de Nível Superior - Brasil (CAPES) - Finance Code 001, CNPq and CAPES COFECUB (19/594).

## 8 Acknowledgments

We thank the CEFAP-USP, especially Dr. Mario Costa Cruz for the confocal imaging and Paulo Albe for preparing the histology slides. This research was supported by Fundação de Amparo à Pesquisa do Estado de Paulo (FAPESP, Grant No 2014/05496-0 and 2017/05264-7). This study was financed in part by the Coordenação de Aperfeiçoamento de Pessoal de Nível Superior - Brasil (CAPES) - Finance Code 001, CNPq and CAPES COFECUB (19/594).

## 1 Data Availability Statement

All relevant data is contained within the article: The original contributions presented in the study are included in the article/supplementary material, further inquiries can be directed to the corresponding author/s.

Datasets are available on request: The raw data supporting the conclusion of this article will be made available by authors, without undue reservation.

## Notes

https://www.ncbi.nlm.nih.gov/bioproject/944921

### Competing Interest Statement

The authors have declared no competing interest.

