## Supplementary figures for "Lack of mTORC2 signaling in CD11c^+^ myeloid cells inhibits their migration and ameliorates experimental colitis"

### 1 Supplementary Figures and Tables

#### 1.1 Supplementary Figures

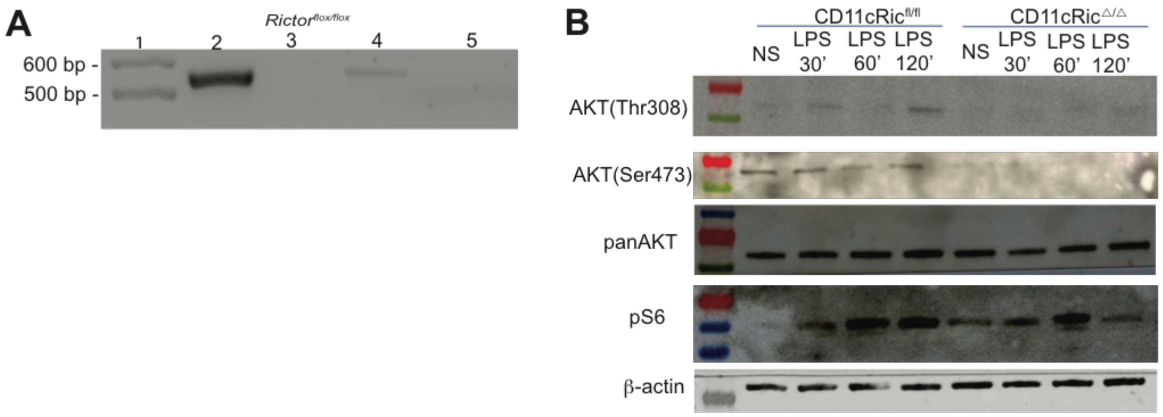


**Supplementary Figure 1. Analysis of mTORC2 signaling ablation in CD11c$^+$ cells and its effect during colitis**. **A,** *Rictor* excision were evaluated on CD11c$^+$ myeloid cells isolated from spleen from CD11cRic$^{fl/fl}$ and CD11cRic$^{\Delta/\Delta}$ (n = 2) mice. Lane 1 = 1Kb DNA ladder, lane 2 = splenic CD11c$^+$ cells DNA from CD11cRic$^{fl/fl}$, lane 3 = splenic CD11c$^+$ cells DNA from CD11cRic$^{\Delta/\Delta}$, 4 = tail DNA from CD11cRic$^{fl/fl}$ (positive control), lane 5 = non-template control (water). **B**, Bone marrow-derived CD11c$^+$ myeloid cells from CD11cRic$^{fl/fl}$ and CD11cRic$^{\Delta/\Delta}$ (n = 2) were generated and expression of mTOR pathway proteins were analyzed. Data representative of 2 independent experiments. **c,** Representative histology of proximal, middle and distal colon sections from CD11cRic$^{fl/fl}$ (n = 5-7)

and CD11cRic$^{\Delta/\Delta}$ (n = 8-10) after treatment with DSS 3% for 7 days. Data representative of two independent experiments.

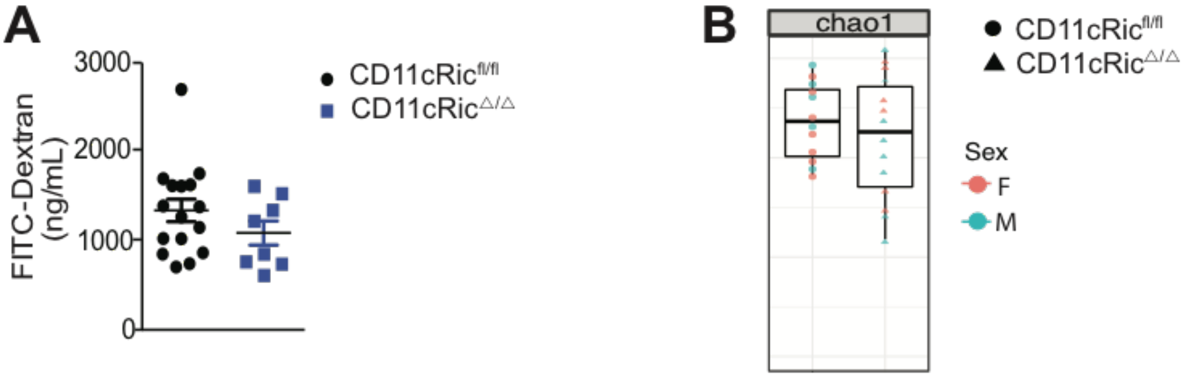


**Supplementary Figure 2. Intestinal permeability and microbiota composition in CD11c Rictor-deficient mice. a**, Intestinal permeability evaluated by FITC-dextran in the serum from CD11cRic$^{fl/fl}$ (n = 13) and CD11cRic$^{\Delta/\Delta}$ (n = 9) mice after treatment with DSS 3% for 7 days. Data is represented as mean ± SEM from pool of two independent experiments. **b**, Alpha diversity analysis of microbiota composition prior DSS-induced colitis. CD11cRic$^{fl/fl}$ (n = 11) and CD11cRic$^{\Delta/\Delta}$ (n = 15) pooled of two independent experiments.

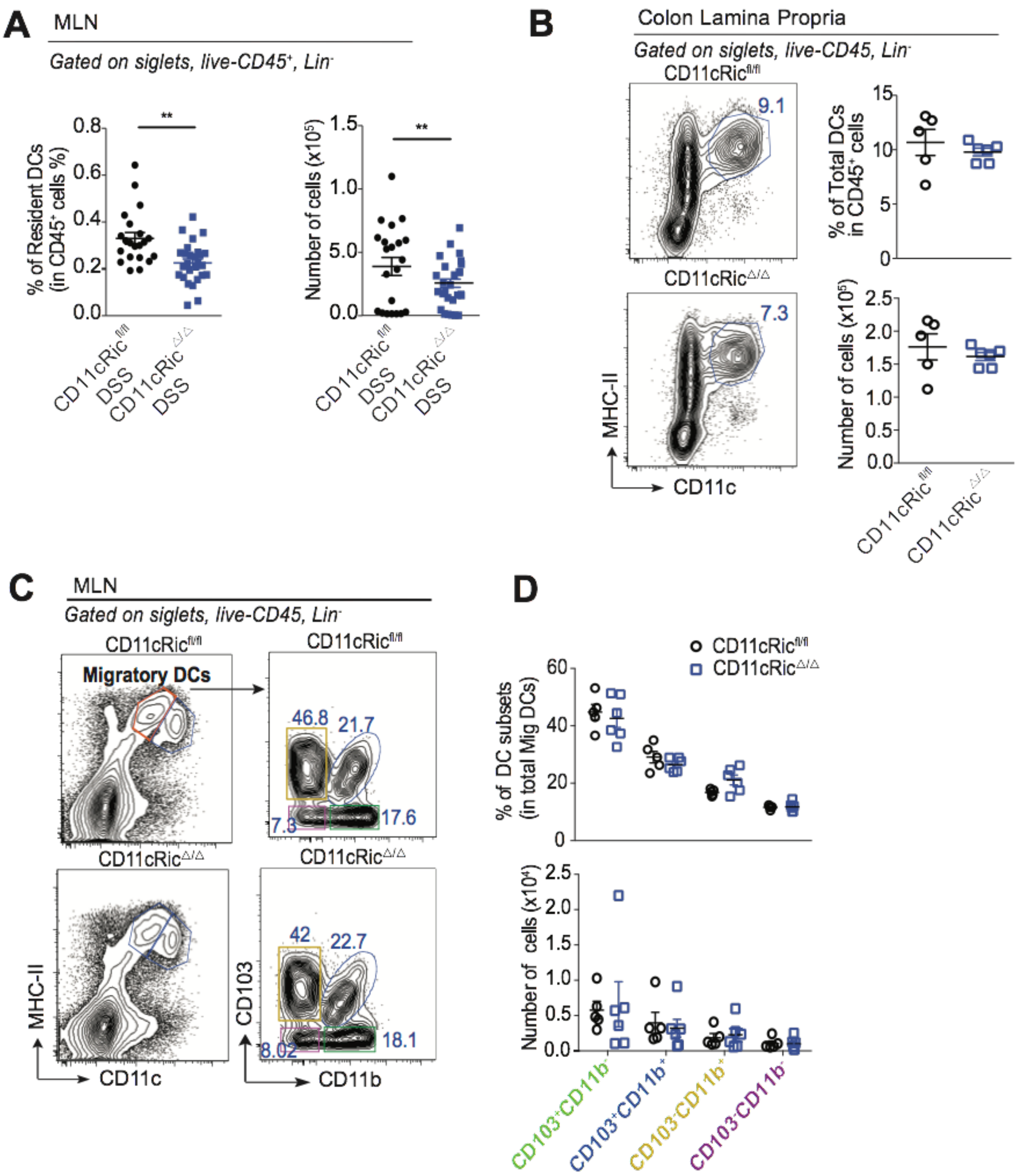


**Supplementary Figure 3. Lack of mTORC2 signaling in CD11c$^+$ cells impacts their abundance during colitis and not under homeostasis. A**, Mice were treated with DSS 3% in the drinking water for 7 days. DCs in the resident compartment of the MLN are shown. CD11cRic$^{fl/fl}$ (n = 7 - 8/experiment) and CD11cRic$^{\Delta/\Delta}$ (n = 8 - 13/experiment). Student t test $^{**}p < 0.01$. Data are expressed as mean ± SEM of three independent experiments pooled. **B-D**, DC analysis during homeostasis. **B**, Total DCs on cLP. **C-D**, DCs subsets on MLN. (n = 2-3/ experiment). Data are expressed as mean ±

SEM of two independent experiments pooled. DCs in the cLP were identified as singlets, live, CD45$^{+}$, Lin$^{-}$ (CD3$^{-}$, B220$^{-}$, Ly6G$^{-}$, SiglecF$^{-}$, Nkp46$^{-}$, CD64$^{-}$) CD11c$^{+}$MCHII$^{+}$. MLN DCs were identified as singlets, live, CD45$^{+}$, Lin$^{-}$ (CD3$^{-}$, B220$^{-}$, Ly6G$^{-}$, SiglecF$^{-}$, Nkp46$^{-}$, CD64$^{-}$) CD11c$^{+}$MCHII$^{low}$ (resident) and CD11c$^{+}$MCHII$^{high}$ (migratory).

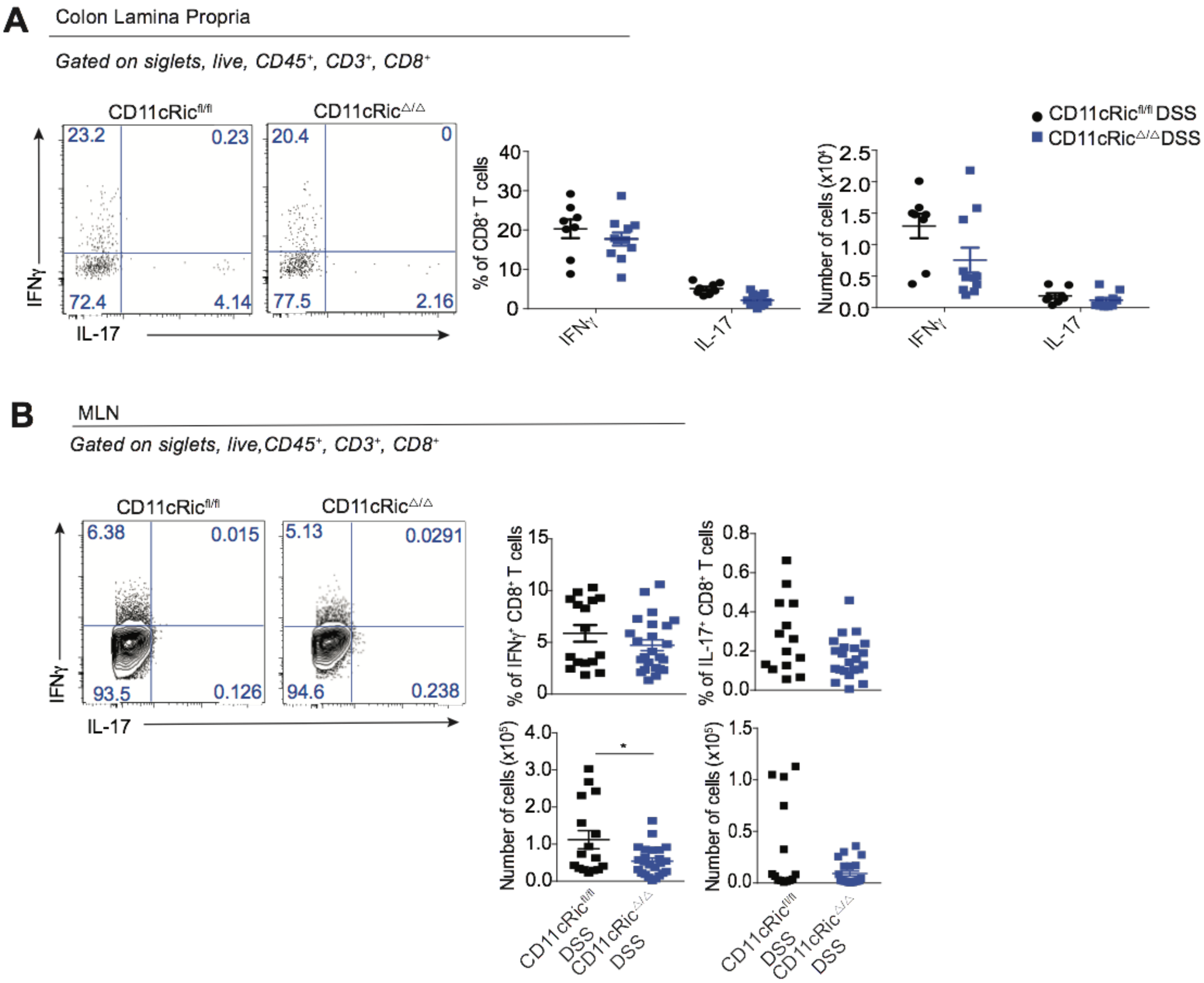


**Supplementary Figure 4. Analysis of CD8$^{+}$ T cells post DSS-induced colitis.** Mice were treated with DSS 3% in the drinking water for 7 days. **A**, IFN-$\gamma$ and IL-17 production on CD8$^{+}$ T cell from cLP. **B**, IFN-$\gamma$ and IL-17 production on CD8$^{+}$ T cell from MLN. CD11cRic$^{fl/fl}$ (n = 7 -8 /experiment) and CD11cRic$^{\Delta/\Delta}$ (n=5 - 6/experiment). Student t test *p < 0.05, **p < 0.01, ***p < 0.001. Data are expressed as mean ± SEM from two independent experiments pooled.

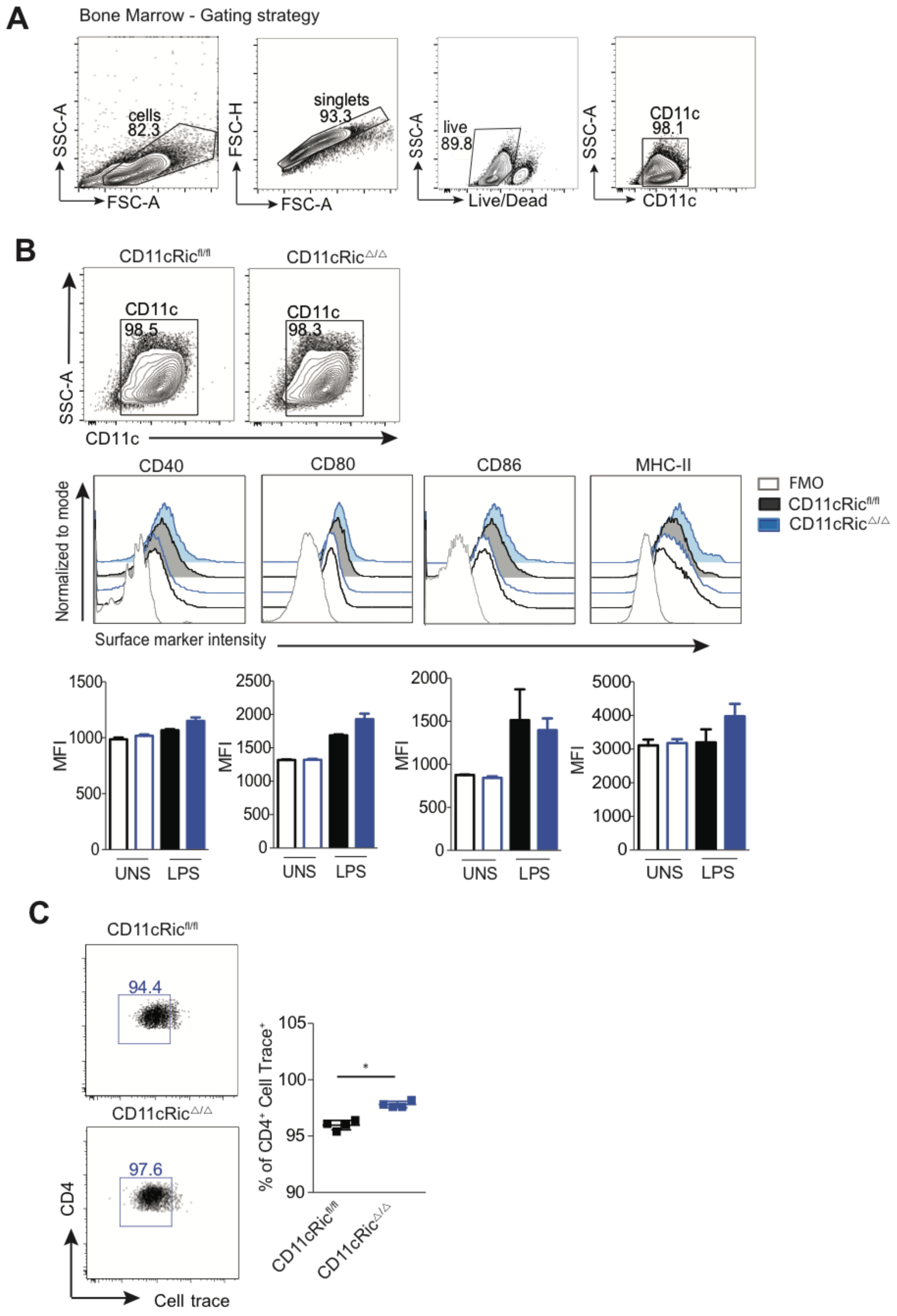


**Supplementary Figure 5. Lack of mTORC2 in CD11c$^+$ compartment impacts their cellular function.** **a**, Gating strategy of CD11c$^+$ cells purified from GM-CSF bone marrow-derived myeloid cells culture. **b**, Analysis of costimulatory molecules on CD11c$^+$ cells stimulated or not with LPS (100 ng/mL) for 24 h. **c**, OVA-pulsed CD11c$^+$ cells were co-cultivated with naïve OT-II CD4$^+$ T cells for 5 days and proliferation analysis was performed with Cell Trace. Student t test $*p < 0.05$. Data are expressed as mean ± SEM and representative from two or three independent experiments.
